# Efficient synergistic single-cell genome assembly

**DOI:** 10.1101/002972

**Authors:** Narjes S. Movahedi, Zeinab Taghavi, Mallory Embree, Harish Nagarajan, Karsten Zengler, Hamidreza Chitsaz

## Abstract

As the vast majority of all microbes are unculturable, single-cell sequencing has become a significant method to gain insight into microbial physiology. Single-cell sequencing methods, currently powered by multiple displacement genome amplification (MDA), have passed important milestones such as finishing and closing the genome of a prokaryote. However, the quality and reliability of genome assemblies from single cells are still unsatisfactory due to uneven coverage depth and the absence of scattered chunks of the genome in the final collection of reads caused by MDA bias. In this work, our new algorithm Hybrid *De novo* Assembler (HyDA) demonstrates the power of co-assembly of multiple single-cell genomic data sets through significant improvement of the assembly quality in terms of predicted functional elements and length statistics. Co-assemblies contain significantly more base pairs and protein coding genes, cover more subsystems, and consist of longer contigs compared to individual assemblies by the same algorithm as well as state-of-the-art single-cell assemblers SPAdes and IDBA-UD. Hybrid *De novo* Assembler (HyDA) is also able to avoid chimeric assemblies by detecting and separating shared and exclusive pieces of sequence for input data sets. By replacing one deep single-cell sequencing experiment with a few single-cell sequencing experiments of lower depth, the co-assembly method can hedge against the risk of failure and loss of the sample, without significantly increasing sequencing cost. Application of the single-cell co-assembler HyDA to the study of three uncultured members of an alkane-degrading methanogenic community validated the usefulness of the co-assembly concept.

## 1 Introduction

Enormous progress towards ubiquitous DNA sequencing has brought a realm of exciting applications within reach, including genomic analysis at single-cell resolution. Single-cell genome sequencing holds great promise for various areas of biology including environmental biology^21^. In particular, myriad unculturable environmental microorganisms have been studied using single-cell genome sequencing powered by multiple displacement amplification (MDA)^1–5^. Since the majority of microbes to date are unculturable, single-cell sequencing has enabled significant progress in elucidating the genome sequences and metabolic capabilities of these previously inaccessible microorganisms.

Although single-cell sequencing methods have passed important milestones, such as capturing >90% of genes in a prokaryotic cell^6^ or finishing and closing the genome of a prokaryote using MDA^22^, the quality and reliability of genome assemblies from single cells lag behind those of sequencing methods from multi cells due to a bias arising from MDA. The main factors that affect quality are uneven coverage depth and the absence of scattered chunks of the genome in the final collection of reads. Also, the outcome of MDA is widely variable ranging from total loss of the sample and any information therein to nearly complete reconstruction of the genome. In this sense, an MDA-based single-cell sequencing experiment is currently a gamble that can potentially lead to the loss of the sample and sequencing expenses. We demonstrate in this work how to hedge against this risk through sequencing and co-assembly of few single cells. Our method replaces a single-cell deep sequencing experiment with multiple single-cell shallow sequencing experiments, allowing for the acquisition of information about multiple single cells simultaneously.

## 2 Results

### Colored de Bruijn graph

Algorithmic paradigms for fragment assembly, such as overlap-layout-consensus and de Bruijn graph, depend on the characteristics of sequencing reads, particularly read length and error profile. Overlap-layout-consensus is a paradigm that is usually applied to assembly projects using long reads, and the de Bruijn graph is another widely adopted paradigm that is used for short read data sets^15^. A number of consecutive *k*-mers (a sequence of length *k* nucleotides) replace each read in the de Bruijn graph paradigm. Each *k*-mer is represented by a unique vertex. An edge is present between two vertices if there is a read in which the two respective *k*-mers are consecutively overlapping. When there are at least *k* consecutive common bases, reads share a vertex (respectively *k* + 1 common bases for an edge) along which contigs are efficiently constructed.

**Figure 1:**
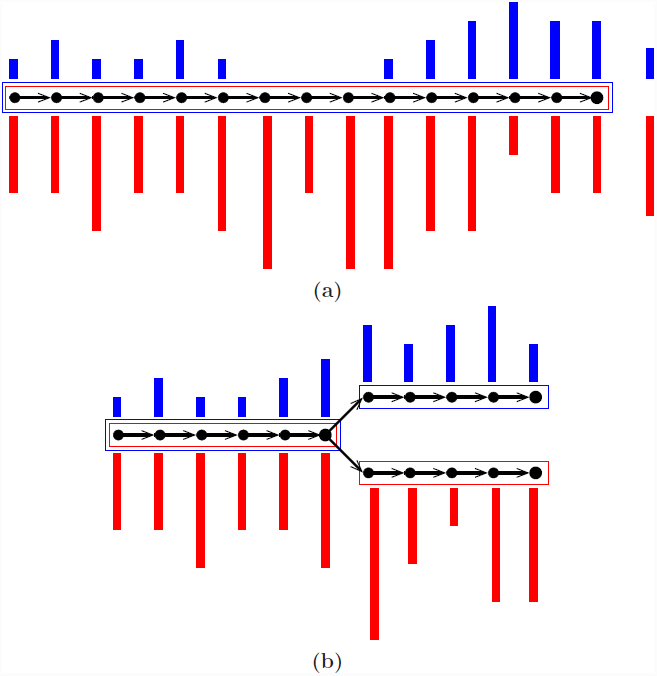
Two sample colored de Bruijn graphs with colors red and blue. Nodes are *k*-mers and edges represent *k* + 1-mers. A colored bar shows multiplicity of the *k*-mer in the corresponding colored input data set. Each box is an output contig, and the color of a box shows non-zero colored average coverage which is shown on the right hand side of the contig in (**a**). Our co-assembly algorithm (**a**) rescues a poorly covered region of the genome in one color when it is well covered in the other, and (**b**) allows pairwise comparison of colored assemblies through revealing all of their shared and exclusive pieces of sequence.

Colored de Bruijn graph is a method proposed for co-assembly of multiple short read data sets^11^. It is an extension of the classical approach by superimposing different uniquely colored input data sets on a single de Bruijn graph. Each vertex, which is a representation of a *k*-mer, accompanies an array of colored multiplicities. In this way, input data sets are virtually combined while they are almost fully tracked, enabling separation after assembly. Iqbal *et al.* proposed the colored de Bruijn graph in Cortex^11^ for variant calling and genotyping, whereas our tool Hybrid *De novo* Assembler (HyDA)^12^ is developed for *de novo* assembly of short read sequences with non-uniform coverage, which is a dominant phenomenon in MDA-based single-cell sequencing^6^. To fill the gaps and compare colors, contigs in HyDA are constructed in a color oblivious manner solely based on the branching structure of the graph. First, this method rescues a poorly covered region of the genome in one data set when it is well covered in at least one of the other input data sets (Figure 1(a), Table 2). Second, it allows comparison of colored assemblies by revealing all shared and exclusive pieces of sequence not shorter than *k* (Figure 1(b), Table 3).

**Figure 2:**
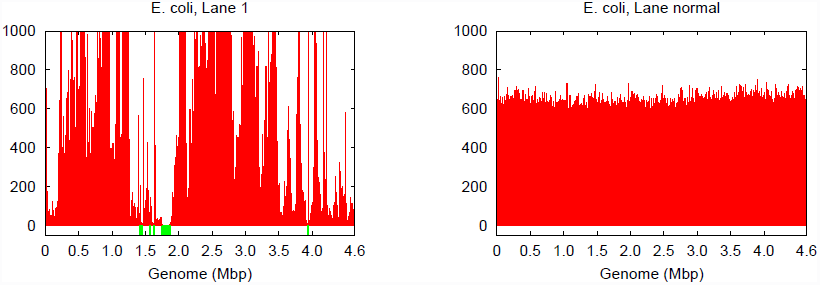
Genome coverage in single-cell *E. coli* lane 1 vs. normal multicell *E. coli*. Both have an average coverage of ~600×.

### Coverage characteristics of single-cell read data sets

Genomes amplified from single cells exhibit highly non-uniform genome coverage and multiple gaps, which are called blackout regions^6^. For the evaluation of such coverage characteristics in this study, we used amplified DNA originating from two single *Escherichia coli* cells as well as from one single *Staphylococcus aureus* cell^6^. Although these amplified DNAs were quality checked for preselected genomic loci using quantitative PCR^8^, they still did not cover the entire genome (Table S1, Figure 2). One single *E. coli* cell was sequenced in four technical replicate lanes (1-4) and the other was sequenced in three technical replicate lanes (6-8) each with a sequencing depth of ~600*×* per lane. The single *S. aureus* cell was sequenced in two technical replicate lanes each with a sequencing depth of ~1,800*×.* All nine lanes were sequenced on Illumina GAIIx platform in paired 2*×*100 bps read mode.

The coverage bias in technical replicates is almost identical, which suggests that the vast majority of bias is caused by MDA. The coverage bias, particularly of the blackout regions, do not always occur at the same genomic loci for different cells of the same genome^6^. Blackout regions in *E. coli* lanes 1 and 6 sequenced from two independently amplified single cells make up 1.8% and 0.1% of the genome respectively, but there are no common blackout regions between these two data sets (Table S1). This means that combining the two data sets could fill all gaps and yield a complete genome, which is the property that HyDA exploits with colored co-assembly.

### Colored co-assembly of *E. coli* and *S. aureus* mitigates the effect of non-uniform coverage

Single-cell read data sets have highly variable coverage^7,8^ (Table S1, Figure 2), which poses serious challenges for downstream applications such as *de novo* assembly. A number of single-cell assemblers including EULER+Velvet-SC^6^, SPAdes^9^, and IDBA-UD^10^ have been developed to mitigate the adverse effects of non-uniform coverage and maximize the transfer of sequencing information into the final assembly. These efforts have been successful, and the existing single-cell assemblers are able to extract nearly all of the information contained in the input data set. However, the vast majority of single-cell data sets do not encompass the entire genome. We report that combining multiple data sets from the same or closely related species significantly improves the final assembly by filling genome gaps (Table S1). The challenge presented by this method is the subsequent deconvolution of single-cell genomes to avoid chimeric assemblies.

The ideal solution involves the co-assembly of multiple data sets without explicitly mixing sequencing reads such that individual assemblies can benefit from the synergy without suffering from chimerism. We propose and implement this solution using the colored de Bruijn graph in HyDA.

We report in Table 2 the co-assembly results for six distinct scenarios (Figure S1), each consisting of a combination of the input read data sets: (i) single-cell assembly of *E. coli* lane 1; (ii) single-cell assembly of *E. coli* lane 6; (iii) mixed monochromatic assembly of *E. coli* lanes 1-4 and 6-8, technical replicates of two biologically replicate single cells; (iv) multichromatic co-assembly of *E. coli* lanes 1-4 and 6-8; (v) mixed monochromatic assembly of non-identical cells: *E. coli* lanes 1-4 and 6-8 and *S. aureus* lanes 7,8; (vi) multichromatic co-assembly of non-identical cells: *E. coli* lanes 1-4 and 6-8 and *S. aureus* lanes 7,8, each assigned a unique color. GAGE, a standard genome evaluation tool, which reports the size statistics and number of substitution, indel, and chimeric errors of an assembly, was used to evaluate our assemblies^18^. In all six scenarios, GAGE results (Table 2) comparing the assembly of color 0 with the *E. coli* reference genome are reported. Color 0 corresponds to *E. coli* lane 1 in (i), (iv), (vi), *E. coli* lane 6 in (ii), and the mixture in (iii), (v) (Figure S1).

**Table 1:** Evaluation results obtained from GAGE^18^ for assembly of *E. coli* lanes 1 and 6 using E+V-SC^6^, SPAdes^9^, and IDBA-UD^10^.

| Tool | Missing Ref. Bases (%) |  |
| --- | --- | --- |
|  | Lane 1 | Lane 6 |
| E+V-SC | 281,060 (6.06%) | 109,994 (2.37%) |
| SPAdes | 128,600 (2.77%) | 15,831 (0.34%) |
| IDBA-UD | 145,536 (3.14%) | 28,583 (0.62%) |

**Table 2:** The GAGE^18^ statistics of HyDA assemblies for the six scenarios in Section 2 (Figure S1). GAGE^18^ is based on MUMmer 3.23 aligner^19.^

|  | Lane 1<br>single color | Lane 6<br>single color | Identical<br>cells<br>mixed | Identical<br>cells<br>colored | Non-identical<br>cells<br>mixed | Non-identical<br>cells<br>colored |
| --- | --- | --- | --- | --- | --- | --- |
| Assembly size | 4,532,221 | 4,642,640 | 5,262,077 | 5,204,061 | 8,273,488 | 5,212,674 |
| Missing *E. coli* | 314,009<br>(6.77%) | 123,687<br>(2.67%) | 1,555<br>(0.03%) | 2,023<br>(0.04%) | 1,289<br>(0.03%) | 2,136<br>(0.05%) |
| Extra bases<br>(%) | 280,998<br>(6.20%) | 198,072<br>(4.27%) | 653,307<br>(12.42%) | 584,534<br>(11.23%) | 3,661,052<br>(44.25%) | 597,088<br>(11.45%) |
| Substitution Error | 60 | 19 | 11 | 3 | 5 | 5 |
| Indels < 5 bp | 6 | 4 | 10 | 6 | 8 | 6 |
| Indels ≥ 5 bp | 13 | 14 | 6 | 5 | 4 | 4 |
| Inversions | 0 | 0 | 0 | 0 | 0 | 0 |
| Relocations | 12 | 11 | 2 | 3 | 2 | 3 |
| NG50 | 42,257 | 54,422 | 41,964 | 34,752 | 54,505 | 37,794 |
| Corrected NG50 | 39,975 | 44,872 | 39,334 | 32,876 | 39,334 | 36,868 |

While the state-of-the-art individual single-cell *E. coli* assemblies by SPAdes (SPAdes outperforms IDBA-UD and Euler+Velvet-SC in this case) miss 128,600 (2.77%) and 15,831 (0.34%) base pairs of the reference genome in the two different single cells (Table 1), our co-assembly misses only 2,023 (0.04%) of the genome (Table 2), an improvement of 126,577 (2.72%) base pairs of the *E. coli* cell 1. Our co-assembly of the two single *E. coli* cells and one *S. aureus* cell misses only 2,136 (0.05%) of the genome. The co-assembly algorithm in this work, without any error correction, *k*-mer incrementation, or scaffolding, increases the total assembly size for both *E. coli* lanes 1 and 6 using only the synergy in the input data sets. Our exclusivity ratio (defined below) obtained from the co-assembly results completely differentiates *E. coli* and *S. aureus* data sets (Table 3).

**Table 3:**
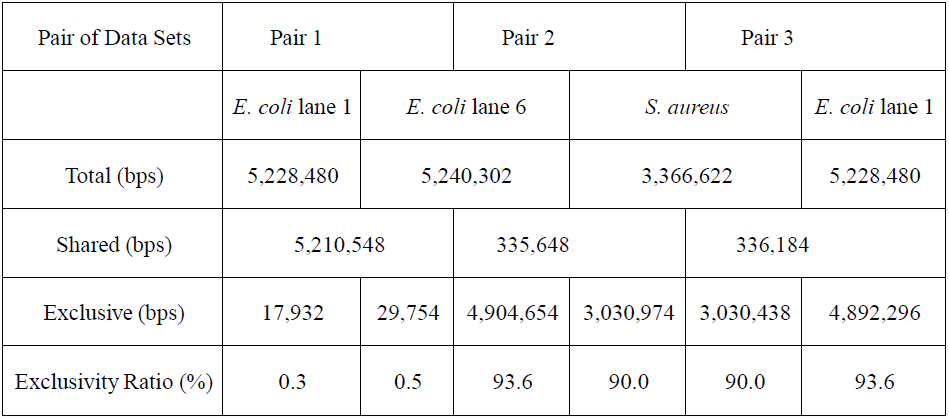
Pairwise relationships between three co-assembled data sets, *E. coli* lanes 1 and 6 and *S. aureus* lane 7, in a co-assembly of *E. coli* lanes 1-4, 6-8 and *S. aureus* lanes 7, 8. Total is the total size of those contigs that have non-zero coverage in the corresponding color. Shared is the size of those contigs that have non-zero coverage in both colors. Exclusive is the size of those contigs that have non-zero coverage in the corresponding color and zero coverage in the other color in the pair. Exclusivity Ratio = Exclusive / Total

### Quantification of similarities and differences between colors

Input data sets can be clustered based on the similarity between their assemblies. For a pair of colors *i* and *j*, contigs belonging to both colors are considered *shared* and contigs belonging to color *i* but not to color *j* are considered *exclusive* of color *i* with respect to color *j*. We define the exclusivity ratio of color *i* with respect to color *j* as the ratio of the size of exclusive color *i* contigs to the total assembly size of color *i*. The exclusivity ratio for *E. coli* lane 1-lane 6 (Pair 1 in Table 3) is less than than 0.5%, while that ratio for *E. coli* and *S. aureus* in the two other pairs (Pair 2 and 3 in Table 3) is greater than 90%. This large difference in exclusivity ratio between Pair 1 and Pairs 2 and 3 is expected in this case, as *E. coli* and *S. aureus* are phylogenetically divergent species belonging to different phyla.

### *De novo* single-cell co-assembly of members of an alkane-degrading methanogenic consortium

The genomes of 10 cells from three dominant but uncultured bacterial members of a methanogenic consortium^13,20^ belonging to the families *Syntrophacea* and *Anaerolineaceae* were sequenced from their amplified single-cell whole DNAs: six cells belonging to *Smithella*, two cells belonging to *Anaerolinea*, and two cells belonging to *Syntrophus*. Single cells were isolated from the consortium by fluorescence-activated cell sorting, and the genomes of individual cells were amplified using MDA. MDA products were sequenced using an Illumina GAIIx with 34, 36, or 58 base pair reads. In total, 10 data sets, one per cell, were obtained.

The 10 data sets were co-assembled with HyDA in a *ten-color* setup, and to exhibit the advantage of the co-assembly method, each data set was assembled individually by HyDA. Individual assemblies created by SPAdes and IDBA-UD were used as comparison. The QUAST^16^ length statistics of the resulting assemblies (≥100 bp contigs) are compared in Table 4 and Figures S2-11. The comparison between individual-assembly and co-assembly by HyDA demonstrates that co-assembly rescues on average 101.4% more total base pairs for all 10 cells (Table S2). Although HyDA does not use advanced assembly features such as variable *k*-mer sizes and paired read information, it can assemble 3.6% to 54% more total base pairs than both SPAdes and IDBA-UD do in all cells except two cases: *Anaerolinea* F02 and *Smithella* MEK03 (Tables 4, S2). When all contigs are considered, HyDA co-assemblies of *Anaerolinea* F02 and *Smithella* MEK03 are 11% smaller and 41% larger than their SPAdes counterparts, respectively. *Smithella* MEK03 input reads are longer (58 bp) than the reads in some of the other data sets; therefore, the *Smithella* MEK03 assembly contains many short contigs and suffers because of the small *k*-mer size (k=25) dictated by the shorter reads.

**Table 4:**
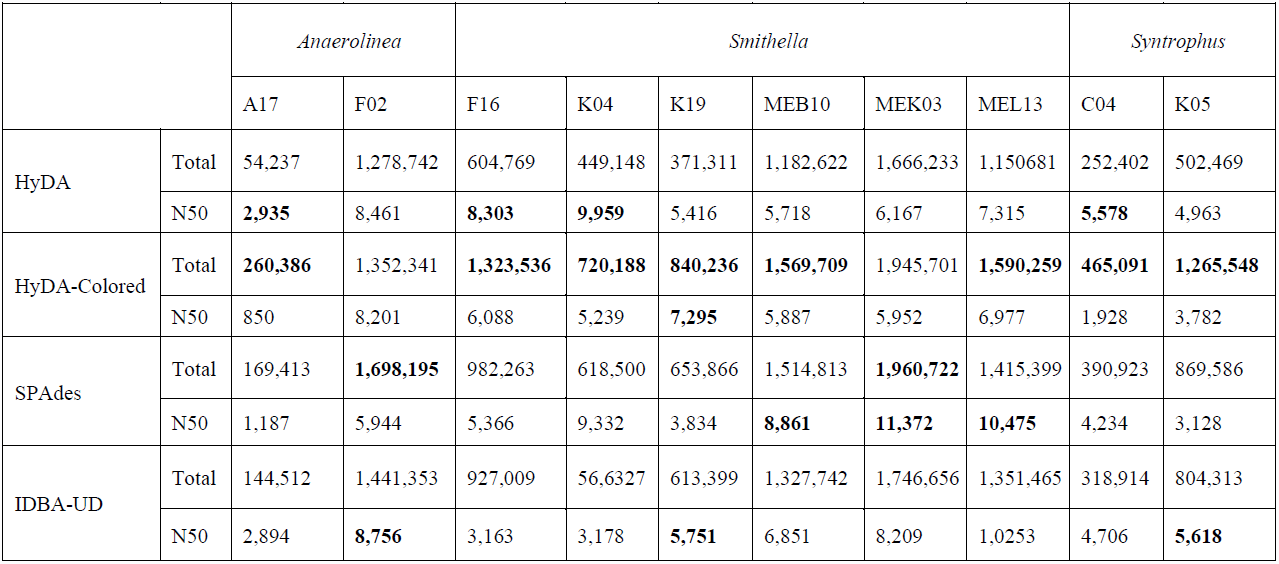
QUAST^16^ analysis of 10 cells from *Anaerolinea*, *Smithella,* and *Syntrophus* single-cell data sets assembled with HyDA (individual assembly), HyDA (10-color co-assembly), SPAdes, and IDBA-UD. All statistics are based on contigs of size >= 100 bp. Only those HyDA contigs that have a coverage of at least 1 in the corresponding color are considered. Coverage cutoff was chosen to be 24 for all HyDA assemblies (-c=24). Total is the total assembly size and N50 is the assembly N50 (the size of the contig, the contigs larger than which cover half of the assembly size).

**Table 5:** The exclusivity ratio (%) of row with respect to column for the 10 cells from *Anaerolinea*, *Smithella* and *Syntrophus* single-cell data sets co-assembled using 10 colors with Squeezambler^17^, a tool in the HyDA package. Only the contigs of coverage at least 1 in the corresponding color are considered. Coverage cutoff was chosen to be 24 for all HyDA assemblies (-c=24).

|  |  | *Anaerolinea* |  | *Smithella* |  |  |  |  |  | *Syntrophus* |  |
| --- | --- | --- | --- | --- | --- | --- | --- | --- | --- | --- | --- |
|  |  | A17 | F02 | F16 | K04 | K19 | MEB10 | Mek03 | MEL13 | C04 | K05 |
| *Anaerolinea* | A17 | 0 | 24 | 87 | 95 | 96 | 80 | 82 | 86 | 22 | 19 |
|  | F02 | 77 | 0 | 96 | 98 | 99 | 95 | 95 | 96 | 74 | 73 |
| *Smithella* | F16 | 96 | 96 | 0 | 73 | 73 | 37 | 22 | 38 | 96 | 55 |
|  | K04 | 97 | 97 | 49 | 0 | 67 | 42 | 25 | 45 | 97 | 57 |
|  | K19 | 98 | 98 | 54 | 68 | 0 | 35 | 32 | 32 | 98 | 58 |
|  | MEB10 | 96 | 96 | 48 | 74 | 69 | 0 | 24 | 39 | 95 | 56 |
|  | MEK03 | 97 | 97 | 49 | 73 | 74 | 38 | 0 | 37 | 96 | 61 |
|  | MEL13 | 97 | 97 | 50 | 76 | 68 | 39 | 22 | 0 | 97 | 59 |
| *Syntrophus* | C04 | 44 | 39 | 89 | 96 | 97 | 85 | 86 | 90 | 0 | 64 |
|  | K05 | 77 | 75 | 54 | 76 | 75 | 45 | 41 | 49 | 73 | 0 |

### Exclusivity analysis of ten assemblies from single uncultured bacterial cells

Exclusivity analysis revealed that the six *Smithella* cells clustered into a consistent group as their exclusivity ratios with respect to the two *Anaerolinea* and two *Syntrophus* cells are almost identical (Table 5). It is important to note that *Anaerolinea* A17 and *Syntrophus* C04 assemblies are relatively short, meaning the exclusivity ratios must be interpreted with caution. Although *Syntrophus* K05’s exclusivity signature with respect to the six *Smithella* cells is indistinguishable from the six *Smithella* signatures with respect to themselves, the exclusivity ratios of *Syntrophus* K05 with respect to the two *Anaerolinea* cells and *Syntrophus* C04 differentiates *Syntrophus* K05 from the six *Smithella* cells. Slight differences between the *Syntrophus* C04 and K05 exclusivity signatures are not surprising because of the existence of potential intraspecies variations.

**Table 6:** Summary of coding sequences and subsystems predicted by the RAST server^14^ for HyDA, IDBA-UD, and SPAdes assemblies of the three alkane-degrading bacterial genomes.

|  |  | HyDA-Colored |  | SPAdes |  | IDBA-UD |  |
| --- | --- | --- | --- | --- | --- | --- | --- |
|  |  | Coding sequence | subsystem | Coding sequence | subsystem | Coding sequence | Subsystem |
| *Anaerolinea* | A17 | **212** | 8 | 146 | **9** | 132 | 7 |
|  | F02 | 1,283 | 122 | **1,653** | **153** | 1,375 | 121 |
| *Smithella* | F16 | **1,197** | **117** | 899 | 91 | 866 | 89 |
|  | K04 | **659** | **89** | 559 | 75 | 508 | 66 |
|  | K19 | **757** | **82** | 581 | 54 | 572 | 57 |
|  | MEB10 | 1,491 | 151 | **1,504** | **156** | 1,297 | 138 |
|  | MEK03 | 1,856 | 180 | **1,955** | **200** | 1,178 | 170 |
|  | MEL13 | **1,535** | **165** | 1,435 | 154 | 1,384 | 148 |
| *Syntrophus* | C04 | **416** | 48 | 375 | **49** | 320 | 36 |
|  | K05 | **1,216** | **121** | 873 | 68 | 854 | 77 |

### Annotation of the *Anaerolinea, Smithella,* and *Syntrophus* assemblies

To assess the quality of co-assemblies with HyDA, IDBA-UD, and SPAdes, we used the RAST server to predict the coding sequences and subsystems present in each assembly. The HyDA assemblies are superior to those of SPAdes and IDBA-UD in terms of the number of coding sequences and captured subsystems for one *Anaerolinea,* four *Smithella,* and both *Syntrophus* assemblies (Table 6). For *Smithella* MEB10 and MEK03, the HyDA assembly closely follows the SPAdes assembly, which provides the largest annotation (Table 6). For *Smithella* F16 and *Syntrophus* K05, HyDA assemblies contain significantly more coding sequences (33% and 39% respectively) and cover more subsystems (29% and 57% respectively) in comparison to the best of SPAdes and IDBA-UD assemblies.

To confirm the accuracy of the assemblies, the closest related species to each assembly was computed by the RAST server. For the HyDA, SPAdes, and IDBA-UD *Anaerolinea* F02 assemblies, the closest species was *Anaerolinea thermophila UNI-1* (GenomeID 926569.3) (no closest genomes data found for *Anaerolinea* A17 by the RAST server). For the HyDA, SPAdes, and IDBA-UD *Smithella* and *Syntrophus* assemblies, the closest species is *Syntrophus aciditrophicus SB* (GenomeIDs 56780.10 and 56780.15). Note that *Syntrophus aciditrophicus SB* is the closest finished genome to the *Smithella* family. This verifies that co-assembly does not create chimeric assemblies, otherwise we would see *Syntrophus aciditrophicus SB* among close neighbors of the *Anaerolinea* assemblies and/or *Anaerolinea thermophila UNI-1* among close neighbors of the *Smithella* and *Syntrophus* assemblies by HyDA.

### Metabolic reconstruction *of Anaerolinea, Smithella,* and *Syntrophus*

Assembly and subsequent annotation of these genomes enables the elucidation of the functional roles of individual, unculturable constituents within the community. *Anaerolinea*, *Syntrophus*, and *Smithella* each represent genera with very few cultured members and only two sequenced genomes—*Anaerolinea thermophila* (no genome paper) and *Syntrophus aciditrophicus*^23^ are the only available sequenced genomes from these genera to date. The only member of *Smithella* that has been isolated, *Smithella propionica*^24^, has not been sequenced yet. In addition to understanding the genetic basis for the unique metabolic capability of this microbial community, the genomes of these particular organisms present an opportunity to explore the breadth of genetic diversity in these elusive genera.

Using the advanced genome assembly algorithm, we recently identified the key genes involved in anaerobic metabolism of hexadecane and long-chain fatty acids, such as palmitate, octadecanoate, and tetradecanoate, in *Smithella*^13^. Based on sequence homology, *Syntrophus* is closely related to *Smithella*, but we cannot determine if it is also actively degrading hexadecane at this point in time.

Only two species of *Anaerolinea* have been isolated and characterized thus far. These species, both isolated from anaerobic sludge reactors, form long, multicellular filaments and are strictly anaerobic^25,26^. Each species is capable of growing on a large number of carbon sources, and both isolates produce acetate, lactate, and hydrogen as the main end products of fermentation. Comparison of the *Anaerolinea sp.* genome derived from single-cell sequencing with the genome of *Anaerolinea thermophila* UN-1 revealed many similarities in potential metabolic capability. The *Anaerolinea* genome obtained from a single cell contains genes for the utilization of galactose and xylose, consistent with a previous physiological characterization of *A. thermophila*^25^. Additionally, the single-cell *Anaerolinea sp.* genome encoded for several transporters and genes related to trehalose biosynthesis, suggesting extended metabolic capabilities of this strain. Furthermore, the genome has an extracellular deoxyribonulease, an enzyme required for catabolism of external DNA, hinting at the strains ability to scavenge deoxyribonucleosides.

## 3 Methods

### Media and Cultivation of the Methanogenic Alkane-Degrading Community

The microbial community was enriched from sediment from a hydrocarbon-contaminated ditch in Bremen, Germany^20^. The consortium was propagated in the laboratory in anoxic medium containing 0.3 g NH_4_Cl, 0.5 g MgSO_4_•7H_2_O, 2.5 g NaHCO_3_, 0.5 g K_2_HPO_4_, 0.05 g KBr, 0.02 g H_3_BO_3_, 0.02 g KI, 0.003 g Na_2_WO_2_•2H_2_O, 0.002 g NiCl_2_•6H_2_O,trace elements and trace minerals as previously described^20^. The medium was sparged with a mixture of N_2_/CO_2_ (80:20 v/v) and the pH was adjusted to 7.0. After autoclaving, anoxic CaCl_2_ (final concentration 0.25 g/L) and filter-sterilized vitamin solution^20^ were added. Cells were supplemented with anoxic hexadecane as previously described^13^. Bottles were degassed as necessary to relieve over-pressurization.

### Single-cell Sorting, MDA, and Genomes Sequencing

Individual cells from the alkane-degrading consortium were obtained by staining (SYTO-9 DNA stain) and sorting of single cells by FACS^13^. Single cells were lysed as previously described and the genomic DNA of individual cells was amplified using whole-genome multiple displacement amplification (MDA)^27^. Amplified genomic DNA was screened for *Smithella-*specific 16S rDNA gene sequences. Six amplified *Smithella* genomes were selected for Next Generation Sequencing. The MDA amplified genomes were prepared for Illumina sequencing using the Nextera kit, version 1 (Illumina) using the Nextera protocol (ver. June 2010) and high molecular weight buffer. Libraries with an average insert size of 400 bp were created for these samples and sequenced using an Illumina Genome Analyzer IIx. 34 bp paired-end reads were generated for K05 (20.9 million reads), C04 (23.3 million reads), F02 (26.9 million reads), and A17 (22.2 million reads). 58 bp single-end reads were generated for MEB10 (41.3 million reads), MEK03 (54.1 million reads), and MEL13 (18.0 million reads). 36bp paired-end reads were generated for F16 (11.0 million reads), K04 (27.2 million reads), and K19 (22.9 million reads).

### Assembly of Single-cell Genomes

Assemblies were obtained using HyDA version 1.1.1, SPAdes version 2.4.0, and IDBA-UD version 1.0.9. SPAdes and IDBA-UD were run with the default parameters in the single end mode. The scripts to generate all of the assemblies are provided in the supplementary material. The length of *k*-mers in the de Bruijn graph was 25, and the coverage cut off to trim erroneous branches in the graph was selected to be 100. The contigs were then annotated using RAST^14^, and the resulting annotation was used to generate a draft metabolic reconstruction using Model SEED^28^. The Whole Genome Shotgun project has been deposited at DDBJ/EMBL/GenBank under the accession AWGX00000000. The version described here is version AWGX01000000.

## 4 Discussion

We demonstrated the power of genome co-assembly of multiple single-cell data sets through significant improvement of the assembly quality in terms of predicted functional elements and length statistics. Co-assemblies without any effort to scaffold or close gaps contain significantly more protein coding genes, subsystems, base pairs, and generally longer contigs compared to individual assemblies by the same algorithm as well as the state-of-the-art single-cell assemblers (SPAdes and IDBA-UD). The new algorithm is also able to avoid chimeric assemblies by detecting and separating shared and exclusive pieces of sequence for input data sets. This suggests that in lieu of single-cell assembly, which can lead to failure and loss of the sample or significantly increase sequencing expenses, the co-assembly method can hedge against that risk. Our single-cell co-assembler HyDA proved the usefulness of the co-assembly concept and permitted the study of three bacteria. The improved assembly gave insight into the metabolic capability of these microorganisms thereby proving a new tool for the study of uncultured microorganisms. Thus, the co-assembler can readily be applied to study genomic content and the metabolic capability of microorganisms and increase our knowledge of the function of cells related to environmental processes as well as human health and disease.

The colored de Bruijn graph uses a single *k*-mer size for all input data sets, which has to be chosen based on the minimum read length across all data sets. For instance, *Smithella* MEK03 input reads are longer (58 bp) than the reads in some of the other data sets, while the *Smithella* MEK03 assembly contains many short contigs because of the small *k*-mer size (k=25) dictated by the shorter reads. This minor disadvantage can be remedied by using advanced assembly features such as variable *k*-mer size, alignment of reads back to the graph and threading, and utilization of paired-end information.

## Acknowledgements

Funding for this work was partially provided by NSF DBI-1262565 grant to H.Ch.

